# The extent of regeneration is impacted by the stage of amputation in ladybird beetles: a case study in *Cheilomenes sexmaculata*

**DOI:** 10.1101/2023.04.20.537621

**Authors:** Hera Alam, Shriza Rai, Praveen C Verma, Geetanjali Mishra

**Affiliations:** Research Scholar, Ladybird Research Laboratory, Department of Zoology, University of Lucknow, Lucknow-226007, UP, India; Principal Scientist, Department of Molecular Biology & Biotechnology, CSIR-National Botanical Research Institute, Lucknow-226001, UP, India; Professor, Ladybird Research Laboratory, Department of Zoology, University of Lucknow, Lucknow-226007, UP, India

**Keywords:** Trade-off, leg length, developmental duration, body weight, resource allocation

## Abstract

Regeneration is a topic of interest for centuries and arthropods are no exception. Trade-offs associated with regeneration are likely to indicate the reallocation of resources from other metabolic activities such as growth, development, or reproduction to regrowth of the lost body part. This may confer negative selection to some of the developmental traits of the organism despite regeneration being overall advantageous. Our hypothesis for the study was that the extent of regeneration and trade-offs associated with it may be stage-specific. To test this hypothesis, the right forelimb of all four larval stages of the ladybird beetle (*Cheilomenes sexmaculata*) was amputated. The amputated stages were reared till adulthood and all developmental transitions were recorded. Leg size was measured and compared with control. The result showed that the regenerated legs of all the treatments were smaller than the control and the regenerative potency of early larval stage was higher than late larval stages. Regeneration causes delay in post-amputation duration in all the treatments, increasing their total development period. However, insignificant difference was observed between the fresh body weight of regenerated and control adults. The length of unamputated left foreleg was reduced in regenerated beetles, showing some internal trade-off.

## 1 Introduction

Regeneration is the restoration of damaged or lost body parts and involves the redevelopment of lost structures into identical or largely similar to the original (Morgan, 1901; Kumar *et al*., 2007; Shah *et al*., 2011). It can be triggered either by autotomy or amputation and can occur at different stages of life-cycle via multiple developmental mechanisms (Seifert *et al*., 2012). First reported in arthropods by French naturalist Reaumur (Dinsmore, 1991), limb regeneration has been investigated across many taxonomic groups (Tanaka, 2003; Tamura *et al*., 2010; Bely and Nyberg, 2010; Agata and Inoue, 2012). Early regeneration studies in arthropods have mainly focused on certain crustaceans (Skinner *et al*., 1985; Hopkins, 1982; Das, 2015) and some hemimetabolous insects such as cockroaches (Truby, 1983; Tanaka *et al*., 1987) and crickets (Lakes and Mücke, 1989; Li *et al*., 2007). The larval morphology of hemimetabolous insects is similar to the adults except for their size (Niwa *et al.,* 2000), thus regeneration occurs during nymphal molts when the epidermis separates from the cuticle (Selin, 2003). However, holometabolous insects differ in limb morphologies between their larval and adult stages (Moran, 1994; Truman and Riddiford, 1999). Also, in holometabolous insects, the pupal stage undergoes repatterning of the whole body, and thereby, non-lethal injuries incurred during larval stages can be repaired during pupation. Thus, the pupal stage provides an ideal opportunity to regenerate the lost structures. For instance, in Coccinellidae *viz. Coccinella septempunctata* L. (Wu *et al*., 2015), *Harmonia axyridis* Pallas (Wang *et al*., 2015; Abdelwahab *et al*., 2018), *Menochilus sexmaculatus* (Fabricius) (Saxena *et al*., 2016) and other ladybirds, limb regeneration occurs during pupation.

The loss of an appendage changes an animal’s morphology which puts constraints on its physiology and biomechanics. This may ultimately have an effect on the performance and behaviour of the injured/amputated organism (Gillis and Higham, 2016). Thus, to survive in a competitive environment, it is required by the animal to regenerate the lost appendage. However, regenerating a structure requires resources that would have been gone to growth and development and can have profound consequences on an organism’s physiology (Maginnis, 2006a). The physiological investment of regrowing a structure also comes with costs, termed as ‘regenerative load’ (Juanes and Smith, 1995) which may not be compensated later in development giving rise to trade-offs.

From insect legs to fish tails to alligator tails (Xu *et al*., 2020), researchers have demonstrated how costly the regeneration process can be (Maiorana, 1977; Bulliere, 1985; Lawrence and Larrain, 1994; Juanes and Smith, 1995). Studies on cockroaches, stick insects and crabs showed that regenerated appendages are complete anatomically, but are smaller than they otherwise would have been (Liangfei *et al.,* 2004; Maginnis, 2006b; Maginnis *et al*., 2014). The reduction in size can vary from slight to extreme and can have negative impact on organism’s foraging, predation, reproduction, survivorship, life span, and dispersal (Salvador *et al*., 1995; Fox and McCoy, 2000; Maginnis, 2006a; Fleming *et al*., 2007, Bateman and Fleming, 2009; Bely 2010; Emberts *et al*., 2019). In some organisms, sensory hairs on regenerated legs are occasionally malformed (Leonovich and Belozerov, 1992; Vollrath, 1995) which impacts the searching rate, reduced mating success and reproductive performance (Uetz *et al*., 1996; Lakes and Mücke, 1989).

Regeneration is the key to adapt in a competing environment. However, to regenerate, the normal development of the organism has to be impacted. Thus, the organism tends to make a trade-off between structural recovery and its associated developmental costs (Michaud *et al*., 2020). With limited resources, several traits which require energy go through allocation conflicts, so that one trait receives more available resources than the other competing trait. To maintain maximal fitness, the organism is required to balance the regeneration with its associated costs. Since this field of trade-offs associated with regeneration have yet not been properly explored, the present study offers some insight into the developmental cost with respect to different larval stages along with the morphology of the regenerated leg.

Under mass rearing in crowded and resource-poor conditions, it is common for coccinellids larvae to cannibalise each other, which imparts a risk of injury to larval cannibals (Schellhorn and Andow, 1999; Michaud, 2003). Around 16 different species of ladybird beetles have been scored on limb regeneration based on their survival, proportion of regenerating leg, and developmental costs associated with limb amputation (Michaud *et al*., 2020). Therefore, ladybirds could be a potential model for regeneration studies and related trade-offs, such as transgenerational (Abdelwahab *et al*., 2018; Michaud *et al*., 2020), developmental (Wu *et al*., 2019a), functional recovery (Yang *et al*., 2016; Wu *et al*., 2019b) and sexual selection (Wang *et al*., 2015). The experiment for regeneration studies was conducted on *Cheilomenes sexmaculata* (Fabricius) (Coleoptera: Coccinellidae). Although the lost appendage during larval stage gets regenerated in the adult (Saxena *et al*., 2016), the regenerated legs were found to be smaller in comparison to the unamputated legs (Unpublished data). To study whether the regenerative potency is related to larval stage, we hypothesized that the trade-off related to regeneration will be specific to the stage of amputation and larvae amputated at earlier stages will have longer leg length compared to those amputated at later stages. Since, regeneration proceeds along the proximo-distal axis (Wu *et al*., 2015), the length of last leg segments will be most affected because of amputation.

## 2 Material and methods

### 2.1 Insect rearing and Stock maintenance

Adults of ladybird beetle, *Cheilomenes sexmaculata,3* (̴ 50 beetles) were collected from local agricultural fields of cowpea plant, *Vigna unguiculata* (L.) from Barabanki, India (26.9268° N, 81.1834° E). Adult beetles were held in 1-liter plastic beakers, filled with aphid-infested plant leaves and twigs, and covered with fine muslin cloth. Beetles were provisioned daily with *Aphis craccivora* Koch (Hemiptera: Aphididae), commonly known as cowpea aphid. Aphids were reared on cowpea plants in polyhouse at 27±2 °C, 65±5% R.H., and 14L:10D photoperiod. Stock culture of beetles was maintained in a controlled room set to 27±2°C, 65±5% R.H. Mated females were removed from beakers and transferred to plastic Petri dishes (9.0×2.0 cm) and provisioned *ad libitum* cowpea aphids. Eggs were collected by transferring females to fresh Petri dish and were held until eclosion. Larvae were kept one per Petri dish and provisioned daily with aphids. Prior to experimentation, beetles were reared for two generations to acclimatize them to laboratory conditions. Egg batches were then randomly selected for experimentation.

### 2.1 Limb amputation of the larval stages

Egg batches collected from stock were randomly divided into five groups of 30 larvae each. Based on the stage of amputation, these groups were assigned to different treatments, *viz.* (1) amputation at first instar larva, (2) amputation at second instar larva, (3) amputation at third instar larva, (4) amputation at fourth instar larva, and (5) no amputation (control). All larvae of the treatment groups were amputated within 24 hours of molting to their specific stages. Prior to amputation, larvae were chilled individually for 5 min and were amputated at proximal end of femur of the right foreleg. Amputation was performed under stereomicroscope (Magnus) with the help of fine micro-scalpel. The amputated larvae were reared in the above mentioned laboratory conditions and were provisioned daily with *ad libitum* aphids, until adult emergence. Larvae were examined daily to record developmental transitions (Instar to instar Developmental Period (DP), Total Developmental Period (TDP), and Pupal Duration (PD)). Post Amputation Developmental Duration (PADD) was recorded for both amputated and unamputated (control) beetles. Fresh body weight (BW) of adults was obtained within 24 hours of adult emergence using an electronic microbalance (Sartorius CP225-D: Sartorius AG, Goettingen, Germany, 0.01mg precision).

### 2.2 Limb size measurement

Upon emergence, right forelegs of adults from each treatments were surgically removed using capsulotomy Vannas scissors and photographed under stereomicroscope (Magnus) at 16x magnification. The contralateral left forelegs of the adult beetles were similarly photographed and were taken as internal controls. The photographs were then measured using ImageJ software (Java 1.8.0_172) (Abramoff *et al*., 2004). The length of right (LLR) and left (LLL) forelegs of adults and their Femur (FeLR & FeLL), Tibia (TiLR & TiLL), and Tarsus (TaLR & TaLL) length were measured.

### 2.3 Statistical Analysis

Data sets were first tested for normality using the Shapiro-Wilk test and were found to be normal for LLR, LLL, TaLL, TDP, DP, PADD & PD and non-normal for FeLR, FeLL, TiLR, TiLL, TaLR & BW. To study the effect of stage of amputation on leg length, leg segment length, different developmental durations, and body weight of adults, data were subjected to General Linear Model (GLM). Treatments were taken as fixed factors and LL, FeL, TiL, TaL, TDP, DP, PAD, PD, and BW were taken as dependent factors. Univariate analysis for variance was performed for TDP, DP, PAD, PD, and BW, whereas for LL, FeL, TiL, TaL multivariate analysis for variance was used to compare left and right leg lengths. The analysis was followed by a comparison of means using post hoc Tukey’s HSD test at α = 0.05. Entire data analyses were performed using SPSS (version 26.0). Leg measurement was performed using ImageJ software (Abramoff *et al*., 2004).

## 3 Results

### 3.1 Effect of amputation stage on leg length

The leg lengths (LLR) of regenerated adults were found to be significantly reduced (except L1) in comparison to control (F= 24.23, df=4, P<0.05). However, leg length of regenerated adult was found to be similar in length with that of control, when amputated at L1 larval stage (Fig. 1). Contralateral left forelegs (unamputated legs, LLL) of regenerated individual of all treatments were significantly smaller in leg length compared to that of contralateral left foreleg of controls (F= 13.10, df=4, P<0.05). Comparison of means revealed insignificant difference between LLR and LLL of respective treatments (Fig. 1).

**Figure 1.**
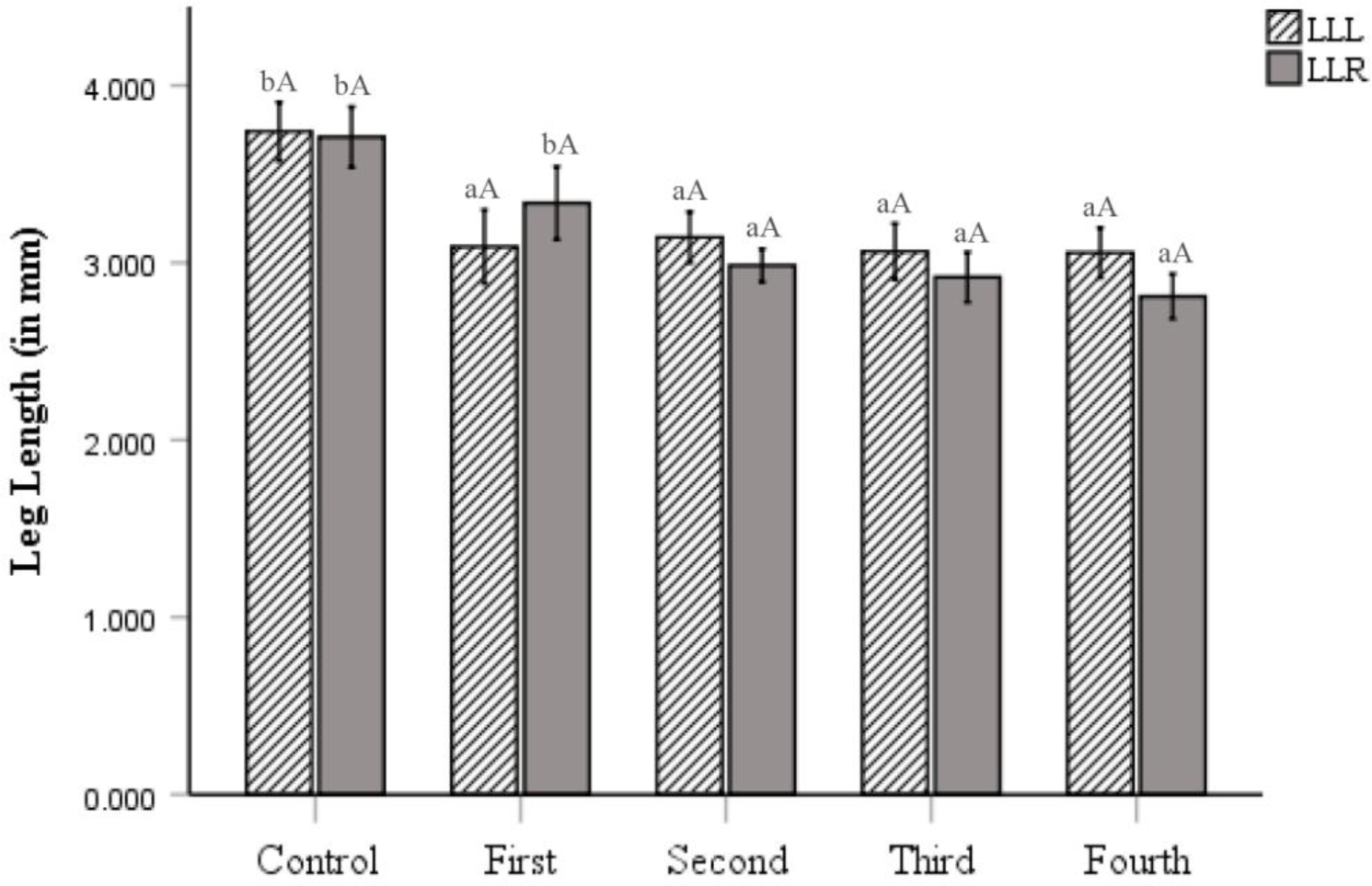
Effect of stage of amputation on leg length of regenerated adults. Values are mean ± SE. Lowercase and uppercase letters indicate the comparison of means across and within the treatments respectively. First, Second, Third and Fourth denotes the larval stage of amputation. Similar letters indicate lack of significant difference at P > 0.05. (LLL=Leg Length Left, LLR = Leg Length Right).

### 3.1 Effect of amputation stage on leg segment length

Depending on the stage of amputation, length of the leg segments (femur, tibia, and tarsus) of regenerated individuals were significantly different from control. All the regenerated leg segments (LR) were found to be significantly affected (F_FeLR_= 13.85, df=4, P<0.05; F_TiLR_= 22.59, df=4, P<0.05; F_TaLR_= 22.10, df=4, P<0.05). Comparison of means amongst the regenerated and control treatments revealed that the femur length (FeLR) of adults amputated at L2, L3, and L4 larval stages was smaller than control. However, insignificant difference was recorded between FeLR of control and L1 amputated adults (Fig. 2a).

**Figure 2.**
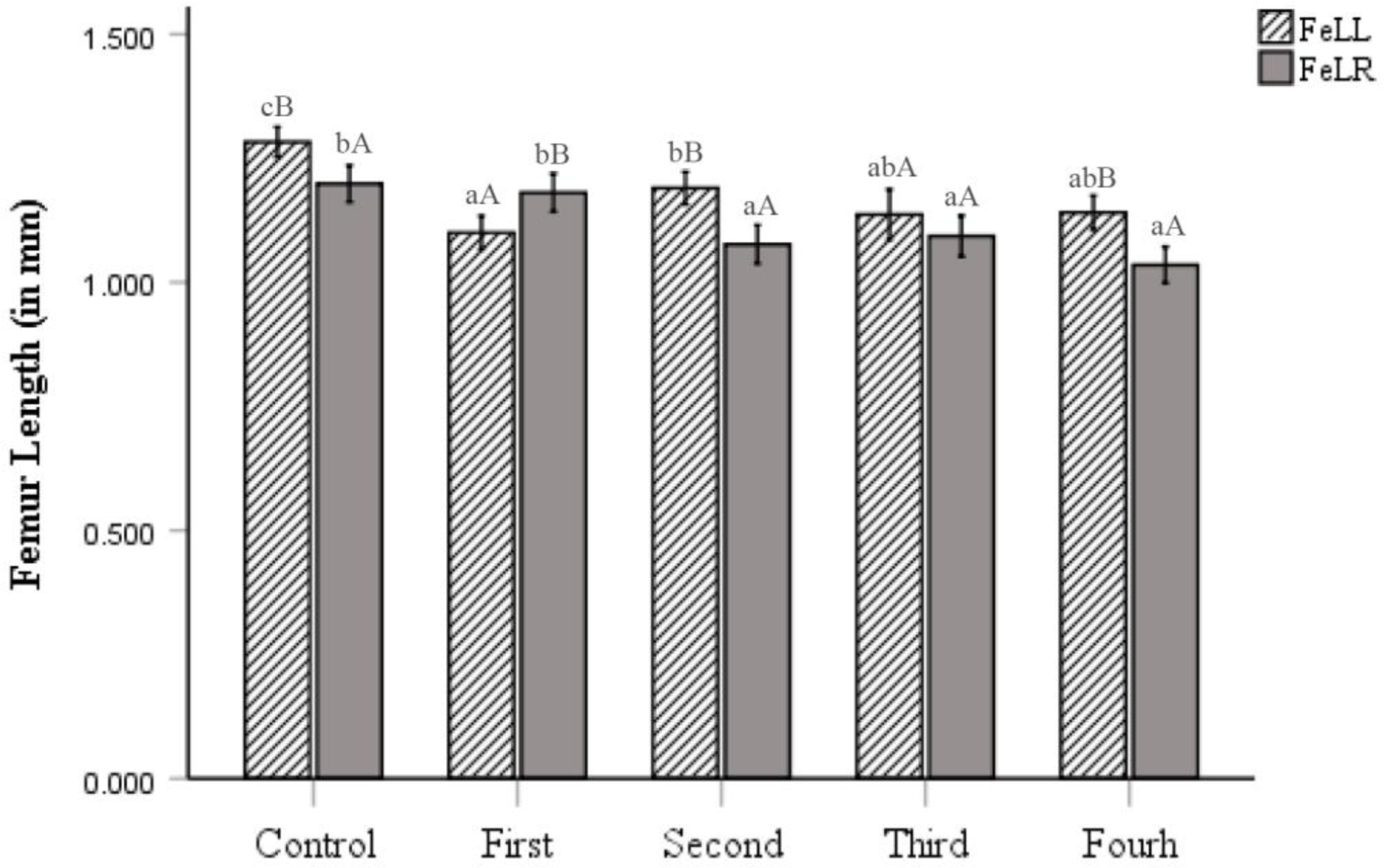
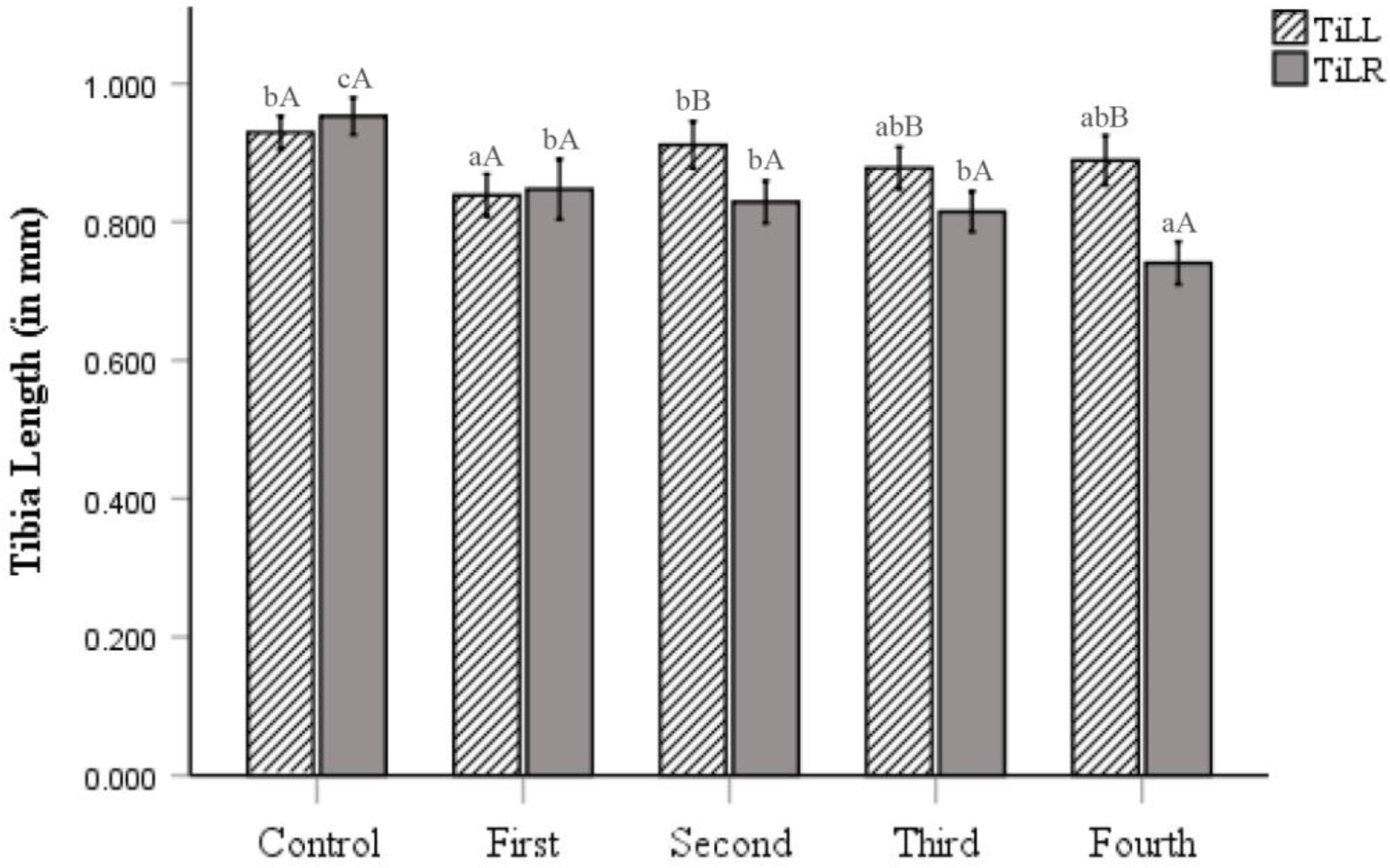
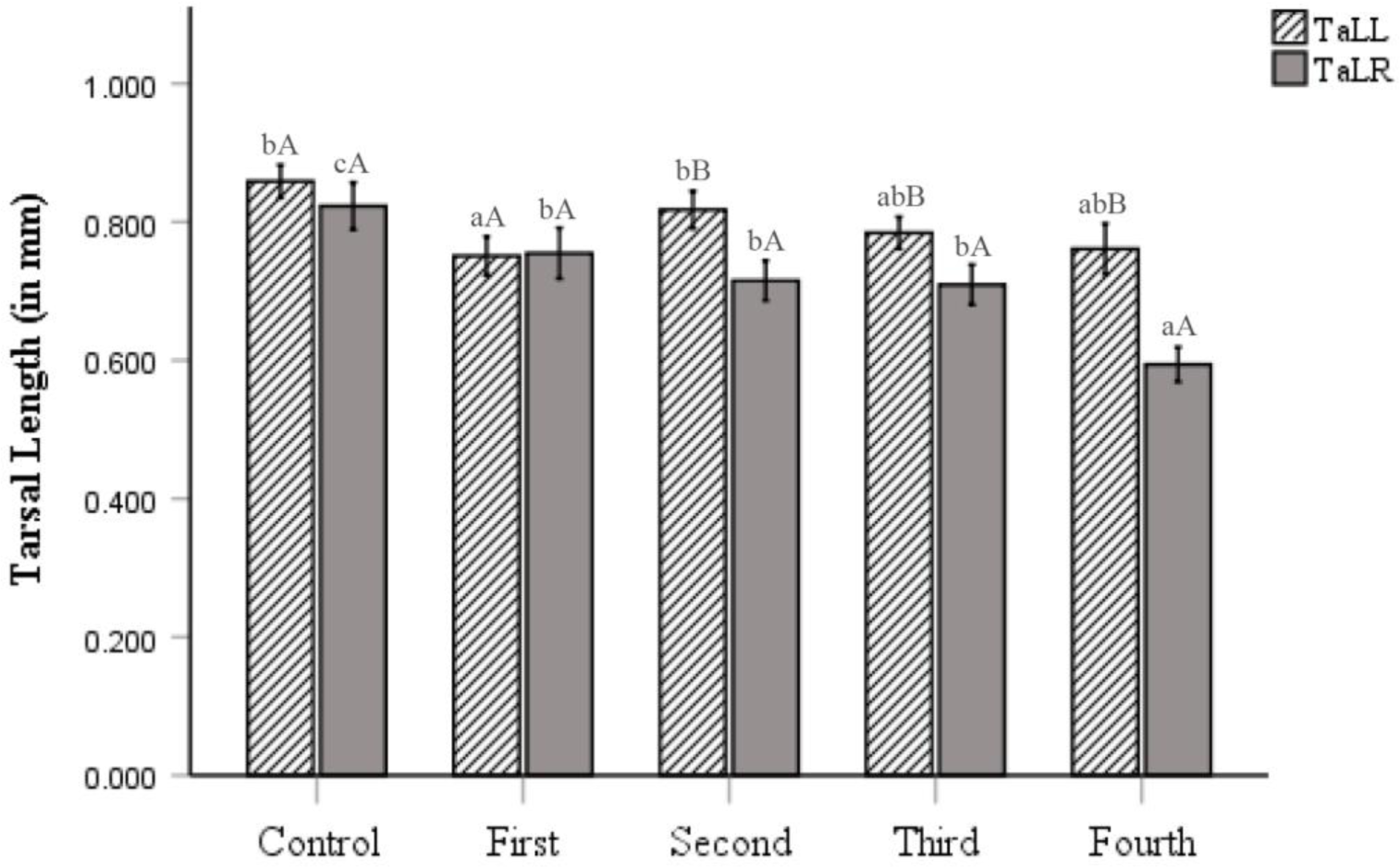
Effect of stage of amputation on leg segment length of regenerated adults. (a) Femur length (b) Tibia length (c) Tarsal length. Values are mean ± SE. Lowercase and uppercase letters indicate the comparison of means across and within the treatments respectively. First, Second, Third and Fourth denotes the larval stage of amputation. Similar letters indicate lack of significant difference at P > 0.05. (FeLL=Femur Length Left, FeLR=Femur Length Right; TiLL=Tibia Length Left, TiLR=Tibia Length Right; TaLL=Tarsal Length Left, TaLR=Tarsal Length Right).

Tibial length (TiLR) of all regenerated legs of adults was found to be smaller than control (F_TiLR_= 22.59, df=4, P<0.05). Comparison of means revealed that the smallest TiLR was observed when larvae were amputated at L4 stage (Fig. 2b). Tarsal length (TaLR) was similarly affected and was smaller in all regenerated adults (Fig. 2c).

Contralateral leg segments (LL) of regenerated adults were also affected (F_FeLL_= 15.14, df=4, P<0.05; F_TiLL_= 5.08, df=4, P<0.05; F_TaLL_= 10.14, df=4, P<0.05). Comparison of means revealed that FeLL of all treatment groups was found to be smaller than control. TiLL and TaLL of L1 amputated adults were smaller than control and no significant difference was found, when the larvae were amputated at L2, L3, and L4 stages.

Comparison within the treatments showed significant difference in the length of leg segments of left (LL) and right (LR) forelegs of adults (Figs. 2a, b, c). FeLR and FeLL of regenerated adults were different when larvae were amputated at L1, L2, and L4 stages. However, the FeLL and FeLR of control were also found to be significantly different. Larvae amputated at L2 and L4 stages had smaller FeLR than FeLL in adults whereas the L1 amputated larvae had smaller FeLL. TiLR of regenerated adults was smaller than TiLL when larvae were amputated at L2, L3, and L4 larval stages. TaLR and TaLL were similarly affected at L2, L3, and L4 larval stages. The tarsal length of right foreleg was smaller than the left foreleg of regenerated adults.

### 3.2 Effect of amputation stage on Total development period (TDP), Development Period (DP), Pupa duration (PD), and Post-Amputation development Duration (PADD)

Total development period (TDP) was observed to be significantly affected by stage of amputation (F=25.89, df=4, P<0.05). Comparison of means revealed that TDP of regenerated larvae was prolonged when the amputation was performed at L1, L2 and L3 larval stages (Fig. 3a). Insignificant difference in TDP was observed between control and L4 amputated larval stages. The developmental period (DP) of larval stages where the leg was amputated was differently affected (F=13.03, df=3, P<0.05). Compared to controls, DP of regenerated L1 larvae was significantly prolonged when the leg was amputated at L1 stage (Fig. 3b). Moreover, the pupal duration (PD) extended significantly in L2, L3, L4 treatments compared to control (F=14.50, df=4, P<0.05) (Fig. 3c). The PADD of all amputation treatments was also found to be significantly higher than the control (F=358.45, df=3, P<0.05) (Fig. 3d).

**Figure 3.**
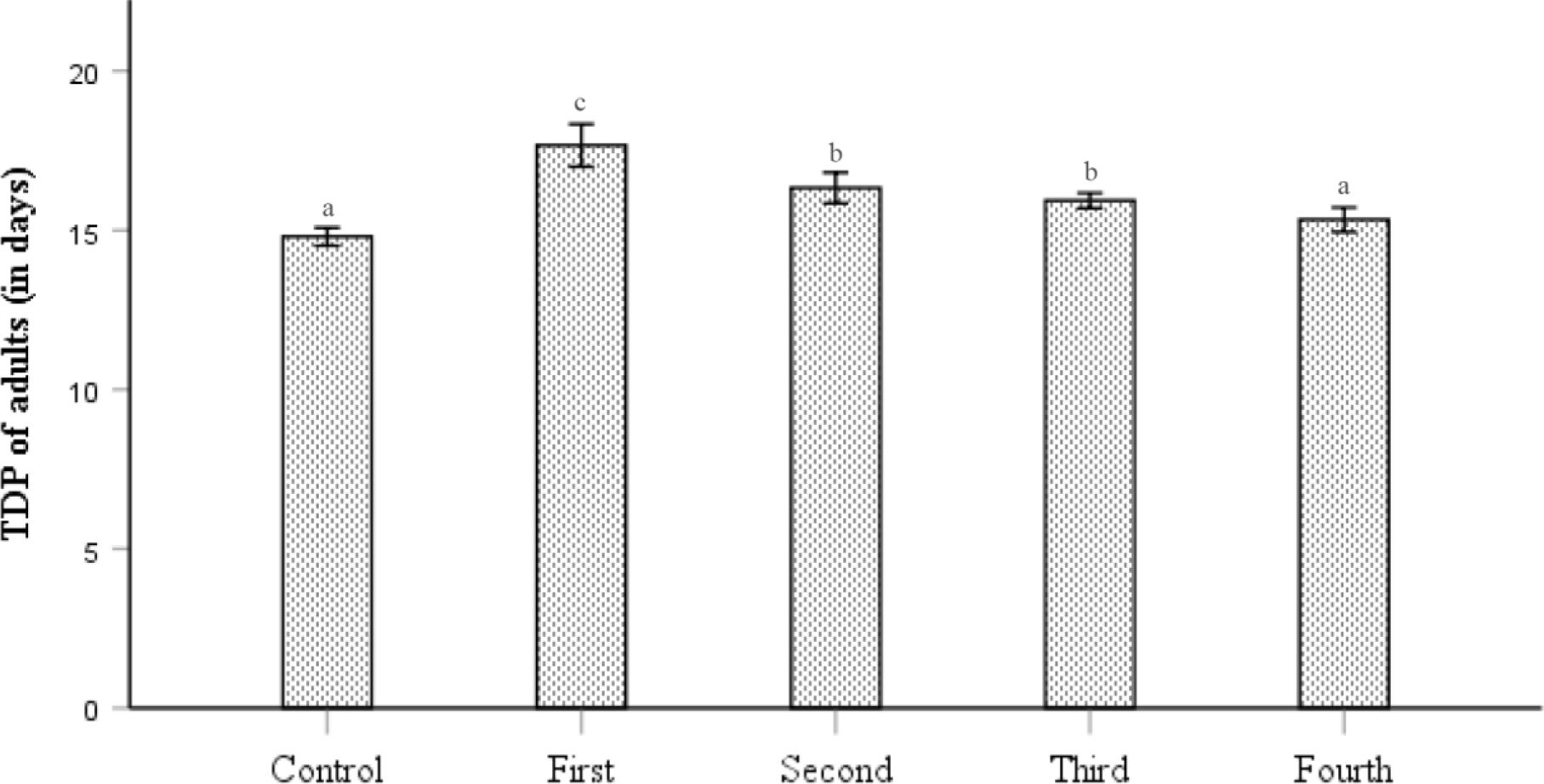
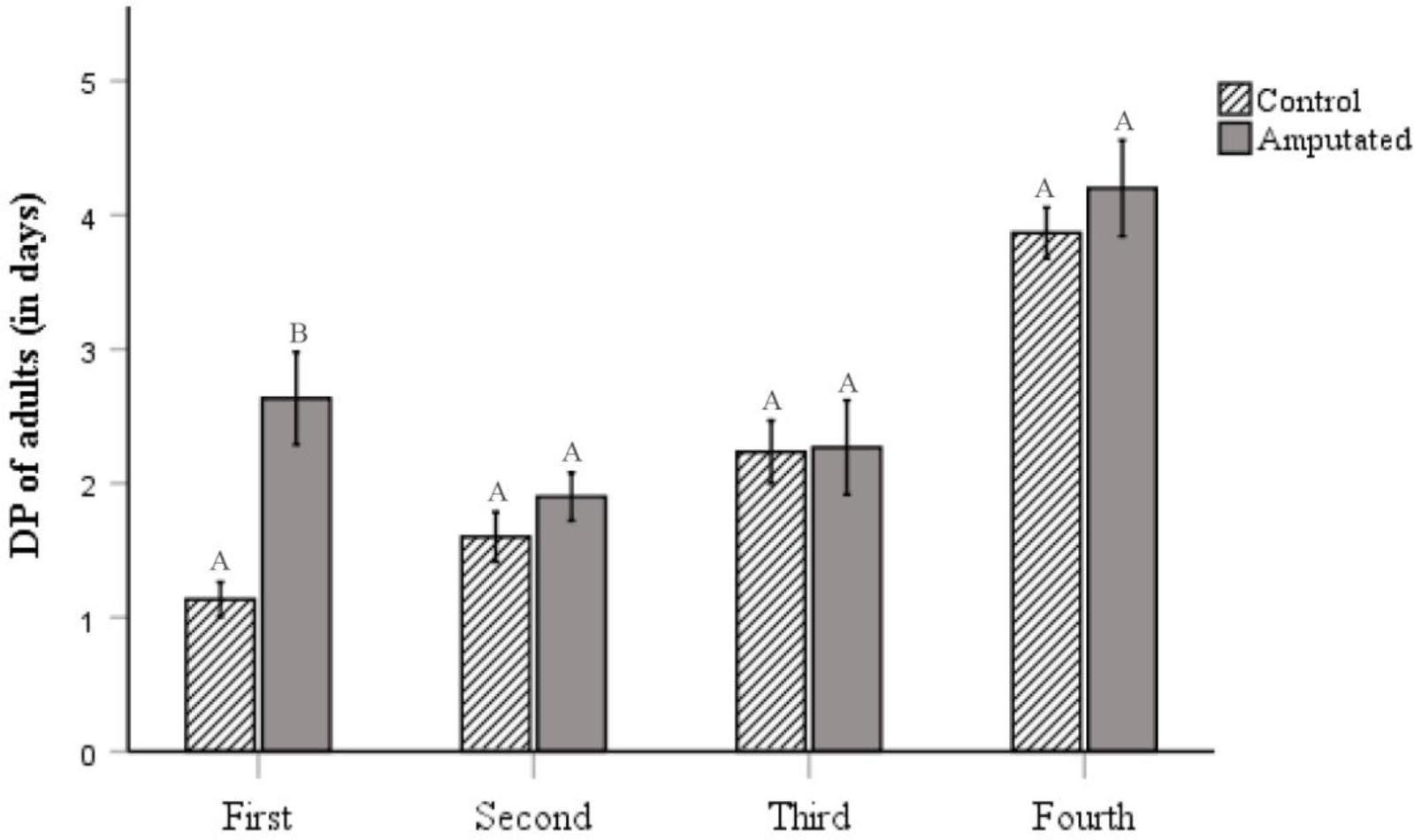
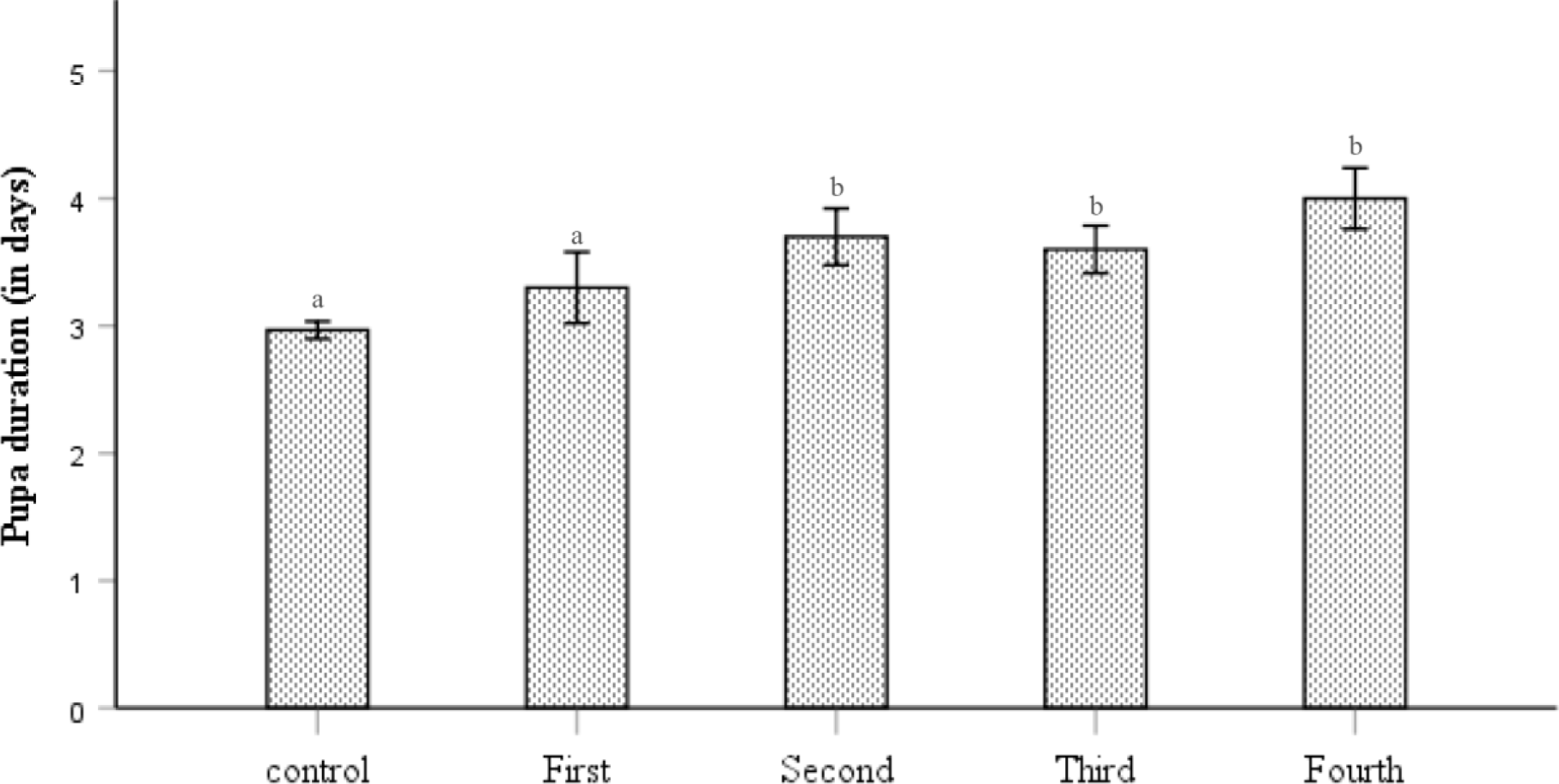
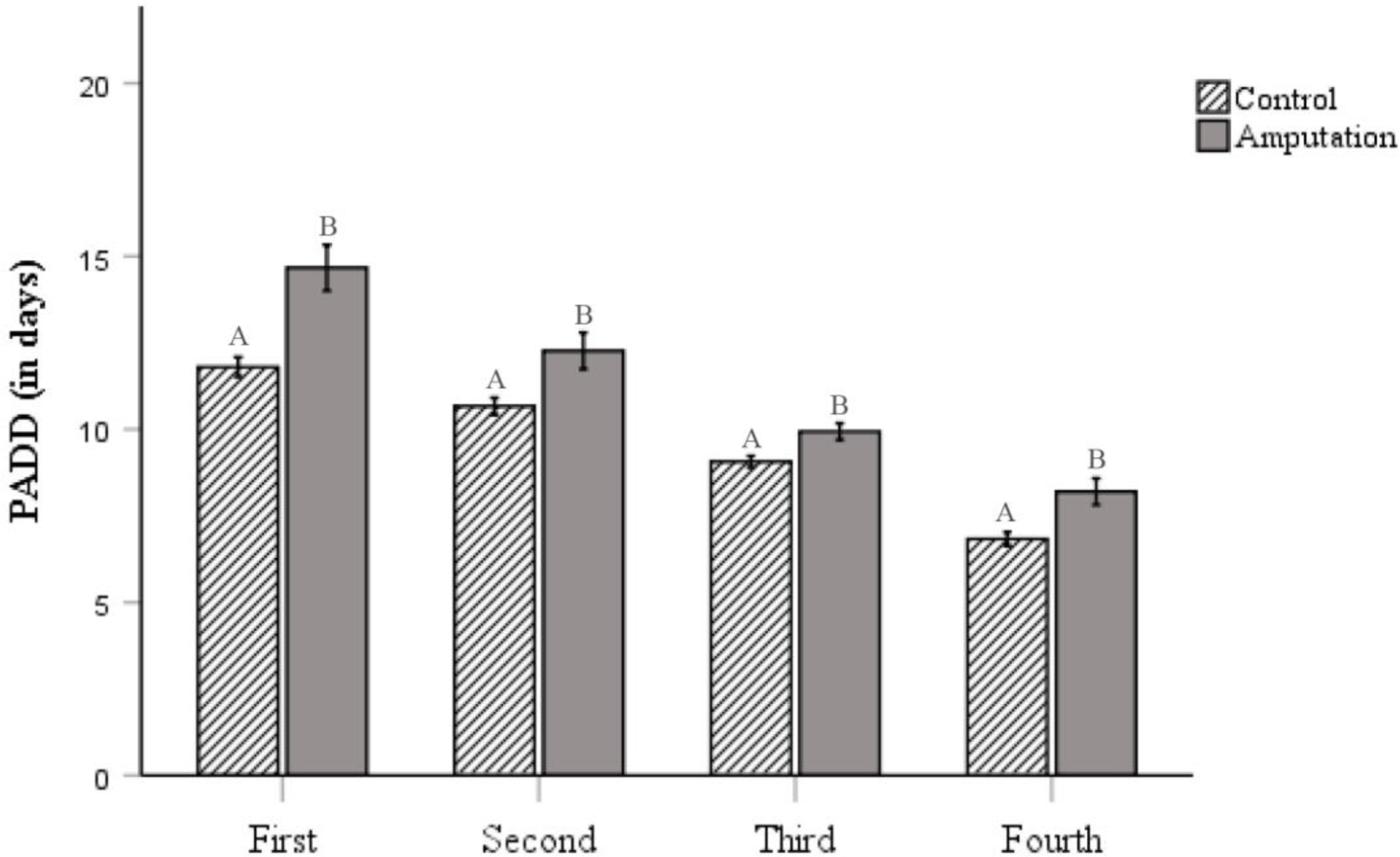
Effect of stage of amputation on different developmental transitions of regenerated adults. (a) TDP (Total development period of adults) (b) DP (development period) of larval stages (c) PD (pupa duration) (d) PADD (post amputation development duration). Values are mean ± SE. Lowercase and uppercase letters indicate the comparison of means across and within the treatments respectively. First, Second, Third and Fourth denotes the larval stage of amputation. Similar letters indicate lack of significant difference at P > 0.05.

### 3.3 Effect of amputation stage on adult Body Weight (BW)

Adult body weight was found to be insignificantly affected by the stage of amputation (F=4.73, df=4 P<0.05). The comparison of mean revealed that there was no significant difference in body weight between adults in the control group and those with amputations at L1, L2, L3, and L4 larval stages (Fig. 4).

**Figure 4.**
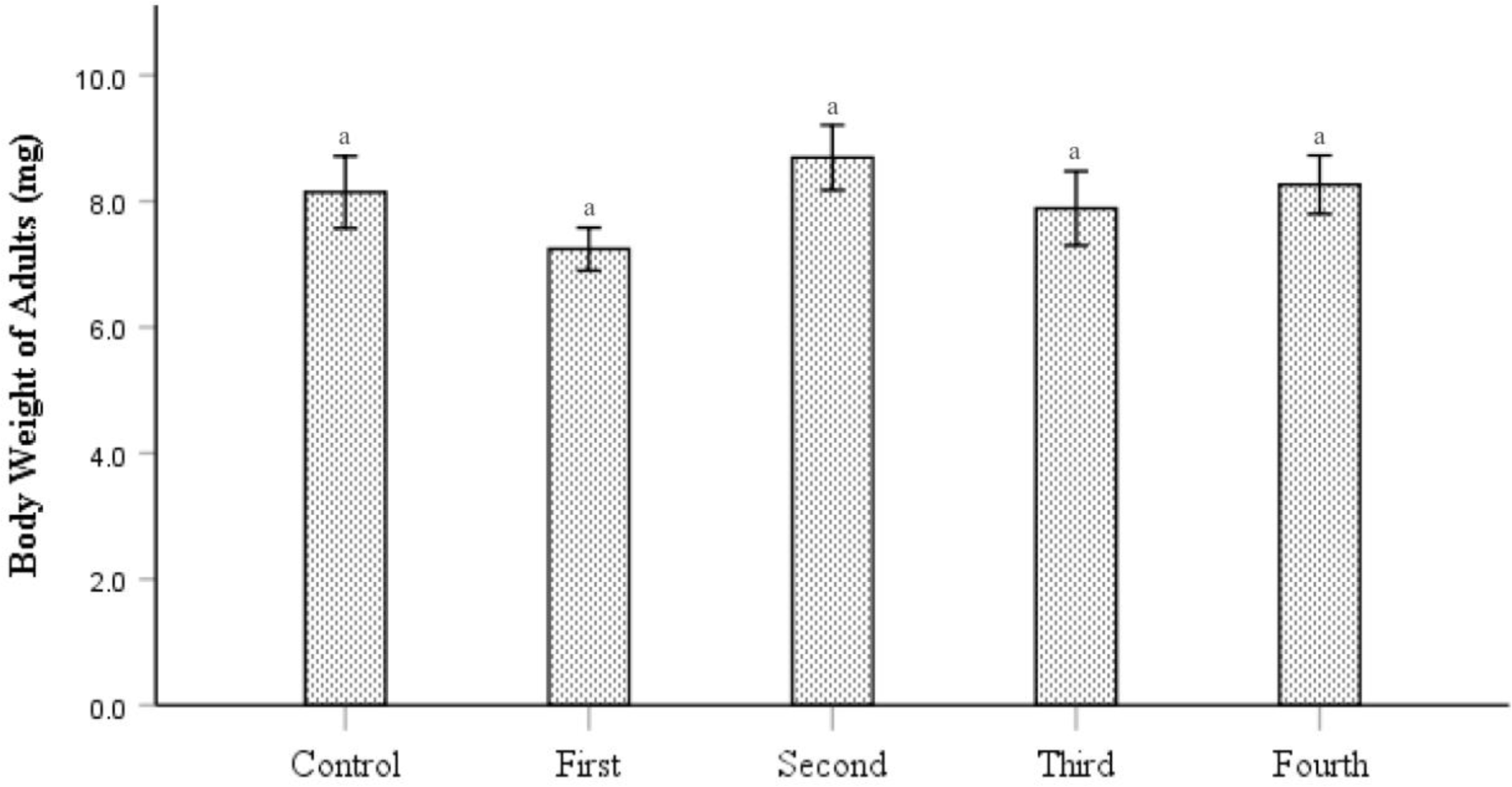
Effect of stage of amputation on BW (body weight) of regenerated adults. Values are mean ± SE. Lower case letters indicate the comparison of means across the treatments. First, Second, Third and Fourth denotes the larval stage of amputation. Similar letters indicate lack of significant difference at P > 0.05.

## 4 Discussion

The present study shows that stage of amputation affects the morphology of the regenerated legs; earlier the amputation better the extent of regeneration. However, the regenerated legs were always smaller than the control.

When the amputation was performed on the first instar larvae, the regenerated legs were of larger size than those with amputation at later larval stages. Hence, the regenerative potency of the early larval stage appears to be higher. The result is consistent with previous studies which show that regenerative ability decreases gradually with increasing larval stages (Roberts, 1973; Shaw and Bryant, 1974; Mengqing and Wanzhi, 2004; Wu *et al.,* 2015). Unlike many arthropods that can regrow limbs after each moult (Maruzzo *et al*., 2005), holometabolous insects are only able to regenerate during the pupal stage. In spite of this, site, location, and stage at which the larva was amputated affect the regeneration potential of adults (Wu *et al*., 2015). This might be attributed to the longer duration of early instar larvae to reach pupal stage, and with each successive moult it might be coping with injury and preparing itself to regenerate during pupal duration.

Not only the amputated foreleg of regenerated adult was affected but its contralateral leg was also shorter in length than those of normal adults. This might be ascribed to the increased utilization of nutritional resources for tissue regeneration following differential allocation of resources from a common resource pool, giving rise to an internal trade-off. Probably because of this reason, use of contralateral leg should be avoided as internal control, earlier taken as normal in regeneration studies by the workers. As the organism tries to maintain its body symmetry, the size of left leg would have been reduced with decrease in the size of right leg. As observed, right and left forelegs show insignificant difference in their size. Regenerated legs were complete anatomically, having all the segments, however, they were smaller than the normal legs. Amputation affected all the segments of regenerated legs in adults with distal leg segments impacted the most in early larval stage (L1). Mainly the distal ends of regenerated legs were curtailed showing regeneration proceeded along the proximo-distal axis (Wu *et al*., 2015). These smaller leg segments affect the morphology of the adult leg which in turn may alter other predatory behaviours, such as prey consumption, searching rate, and handling time in adults (Wu *et al*., 2019b). The reduced size of forelegs is likely to trigger a selected prey acceptance by choosing prey which are smaller and easier to capture. This switch to small-sized suboptimal prey have been observed in many crustaceans, arachnids and insects (Juanes and Smith, 1995; Brueseke *et al*, 2001; Wu *et al*., 2019b). Further study on prey-size preference of the regenerated adults is recommended to comprehend the functionality of the regenerated legs. Although insect and vertebrate legs are different anatomically and have evolved independently, they do show some similarities in their developmental biology. The three classes of secreted factors which control the P/D axis initiation in insects, *i.e.,* Wnt, Hedgehog, TGFβ (Campbell and Tomlinson, 1995) are also found in the developing vertebrate appendage (Parr *et al*., 1993; Riddle *et al*., 1993; Francis *et al*., 1994). Therefore, it is likely that the study of trade-offs in insects may be extended to higher organisms, offering better insights into the evolution of regeneration.

The immediate cost of leg regeneration was the extension in developmental period of the stage at which the larvae were amputated as observed in L1 amputated larvae. The immediate stage was not affected when amputated at L2, L3 and L4 larval instars. However, the latter three stages showed extension in their pupal duration (except for L1 stage). Thus, regenerated adults show increase in their post-amputation development duration (PADD) either by extending their development duration at the stage of amputation or by extending their pupal duration, eventually delaying their TDP (Dewes, 1973; Kunkel, 1977; Wang *et al*., 2015; Wu *et al*., 2019a). The development appears to be ceased and subsequent moults were postponed until the damaged tissue was healed, whereupon development commences and the emerging adult appears to be morphologically normal. Holometabolous insects undergo complete metamorphosis and involve the transformation of the entire body plan during pupal stage. This provides an ideal opportunity to regenerate a lost structure. It is likely that the regenerating tissue triggers a systematic inhibition of development. For instance, in *Drosophila* injured tissue secretes some insulin-like peptides which inhibit ecdysteroid production. As ecdysteroid controls the timing of larval-larval, larval-pupal and pupal-adult transition (Riddiford and Truman, 1993; Yamanaka *et al*., 2013), it causes developmental delay (Poodry and Woods, 1990; Shingleton *et al*., 2005; Garelli *et al*., 2012).

The observed fresh body weight of regenerated adults showed no difference with that of control, indicating no trade-off between regeneration and body weight of the beetles. Probably, the increased PADD in amputated beetles resulted in increased consumption. The extra resource consumption for limb regeneration might come at the expense of other physiological costs. As evident from other studies, allocating resources to regeneration effects both development and reproduction (Dial and Fitzpatrick, 1981; Hill *et al*., 1988; Norman and Jones, 1991; Fleming *et al*., 2007; Bateman and Fleming, 2009; Maginnis *et al*., 2014; Bayoumy *et al*., 2019; Wu *et al*., 2019). In *Coccinella septempuncata* and *Coccinella undecimpunctata,* predatory abilities were reduced following limb regeneration (Bayoumy *et al*., 2019; Wu et al., 2019b). Regeneration led to decrease in wing size, increasing wing loading in stick insect, *Sipyloidea sipylus.* (Maginnis, 2006b). These trade-offs are not only limited to lower organisms but can also be found in higher organisms, such as reduced fecundity observed in skinks (Smyth, 1974), salamanders (Maiorana, 1977), geckos (Dial and Fitzpatrick, 1981), polychaete annelids (Hill et al., 1988) and crabs (Norman and Jones, 1991). Thus, studies focusing on the proximate and ultimate costs of regeneration and their link into population dynamics and ecology will provide a background to explore the evolution of regenerative capacities in animals.

The present study was conducted to observe the morphology of regenerated leg and its associated trade-offs. Studying the trade-offs would perhaps provide more insights as how regeneration may have shaped its presence or absence in the animal kingdom. It will be interesting to assess the impact of regeneration on physiology and morphology of other body parts for future research. Including factors like foraging, locomotion, predation, pattern and speed of regeneration and general life-history traits, into comparisons on an evolutionary level will allow us to specifically look at how regeneration has shaped animal form and function.

## 5 Conclusions

To conclude, the stage of amputation affects the extent of regeneration in adults. Our data provided evidence that the regenerative potency of early larval instars was higher than late larval instars. Regenerated leg was smaller in all cases, even affecting the size of different leg segments viz, femur, tibia, and tarsal length. The contralateral leg (unamputated) of regenerated adults was also smaller, showing some internal trade-off. Based on this finding it is suggested not to use contralateral legs as control. Besides, leg amputation causes extension in post-amputation development duration and delays the total development duration of regenerated adults. It seems, therefore that regeneration utilizes nutritional resources and compromises with the physiology and morphology of the organism.

## 6 Acknowledgements

HA, GM and PV acknowledge SERB-DST (F.No. CRG/2020/001095 via letter no. BP-2020-20-3308 dated December 30, 2020). SR acknowledges UGC Fellowship by University Grants Commission, New Delhi, India (F.No. 16-9(June 2019)/ 2019 (NET/CSIR) dated August 09, 2019).

## 7 Author Contributions

HA: methodology, validation, formal analysis, visualization, writing e original draft, review & editing. SR: methodology, validation, writing e review & editing. PV: writing e review & editing, supervision, validation, funding acquisition. GM: conceptualization, validation, resources, writing e review & editing, supervision, funding acquisition.

